## Supplemental Data for "Regulation of HMGB2 integrates ribosome biogenesis and innate immune responses to DNA"

**List of Supplemental Information**

Supplemental Figures S1 - S5

Tables S1-S4

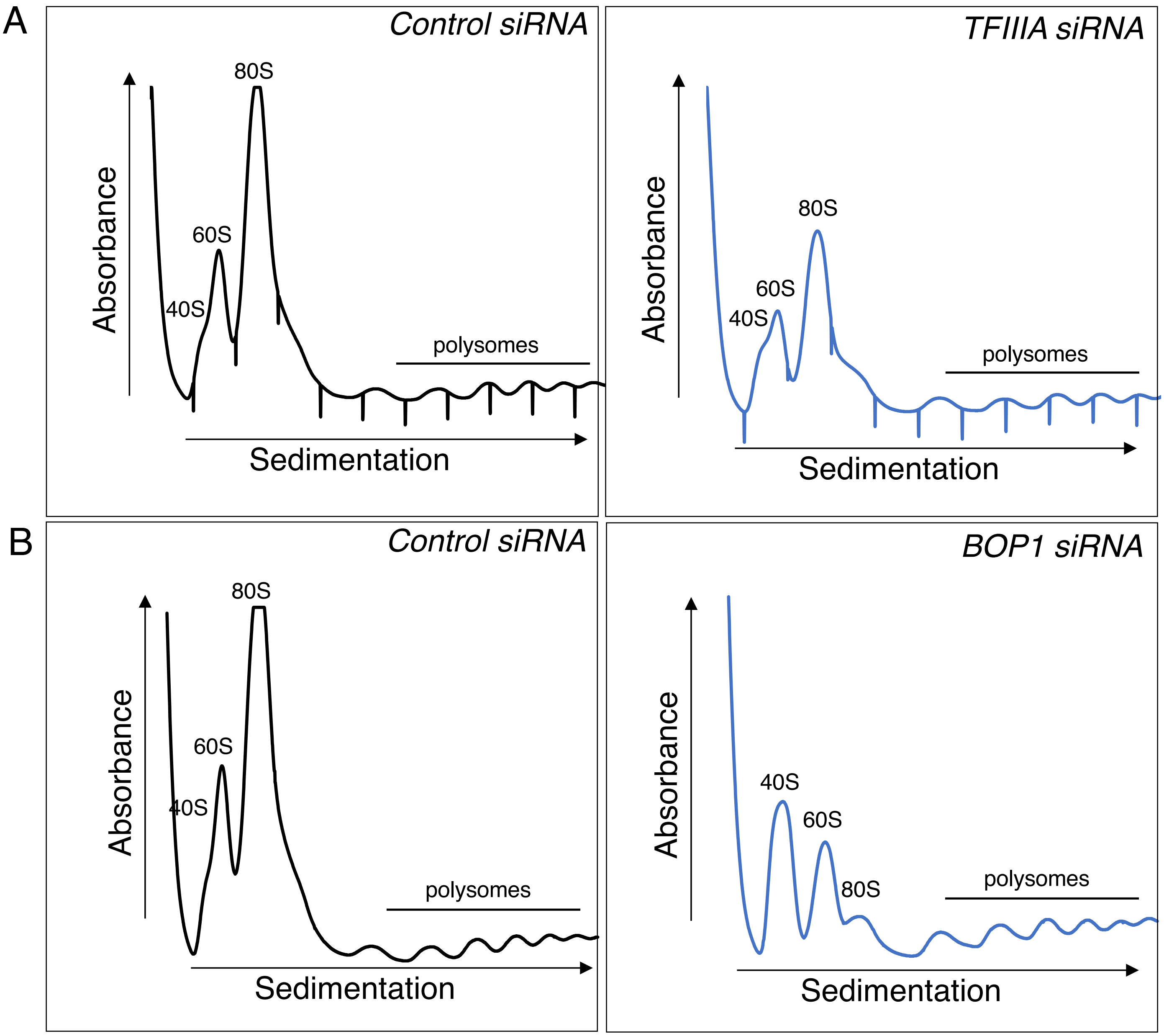


**Figure S1. Depletion of TFIIIA or BOP1 Reduce Ribosome Abundances In HCMV Infected Cells.** (a) NHDFs were transfected with ns or TFIIIA siRNA and infected with HCMV (MOI=3PFU/cell.) At 2 dpi, cytoplasmic lysate was fractionated over a 10-50% sucrose gradient to resolve ribosomal subunits, monosomes, and polysomes, and the absorbance at 254nm was recorded. The top of the gradient is on the left. (b) as in (a) except ns and BOP1 siRNA were used.

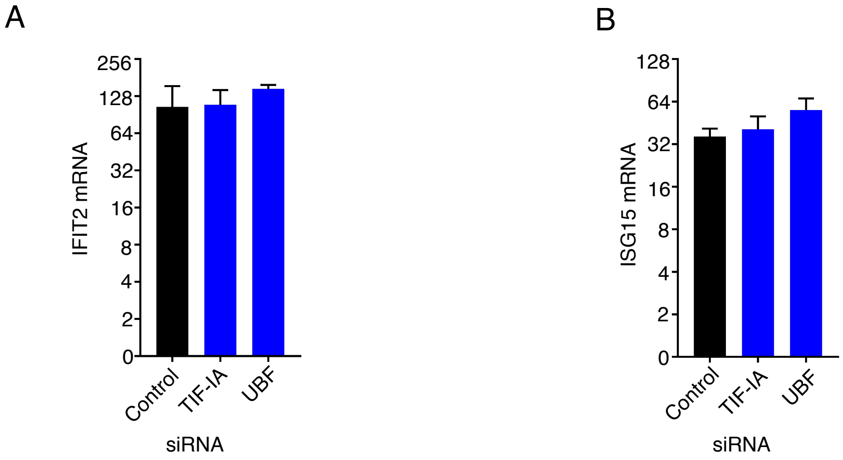


**Figure S2. Depletion of RNA Polymerase I Factors Does Not Impact Interferon Signaling.** (a,b) NHDFs were transfected with ns, TIF-IA, or UBF siRNA. After three days, cells were treated with recombinant IFNα for 6 hours, total RNA was isolated, and RT-qPCR was performed using primers specific for (a) IFIT2 mRNA and (b) ISG15 mRNA.

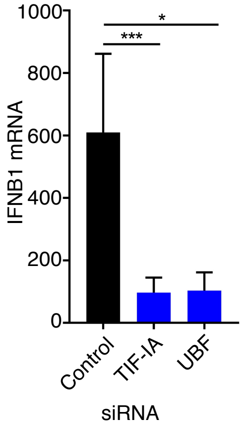


**<u>Figure S3.</u> UBF is Required for Efficient dsDNA-induced accumulation of IFNB1 mRNA.** NHDFs were transfected with ns, TIF-IA, or UBF siRNA. After three days, cells were transfected with dsDNA for 6 h, total RNA was isolated, and RT-qPCR was performed using primers specific for IFNB1 mRNA.

Figure S4

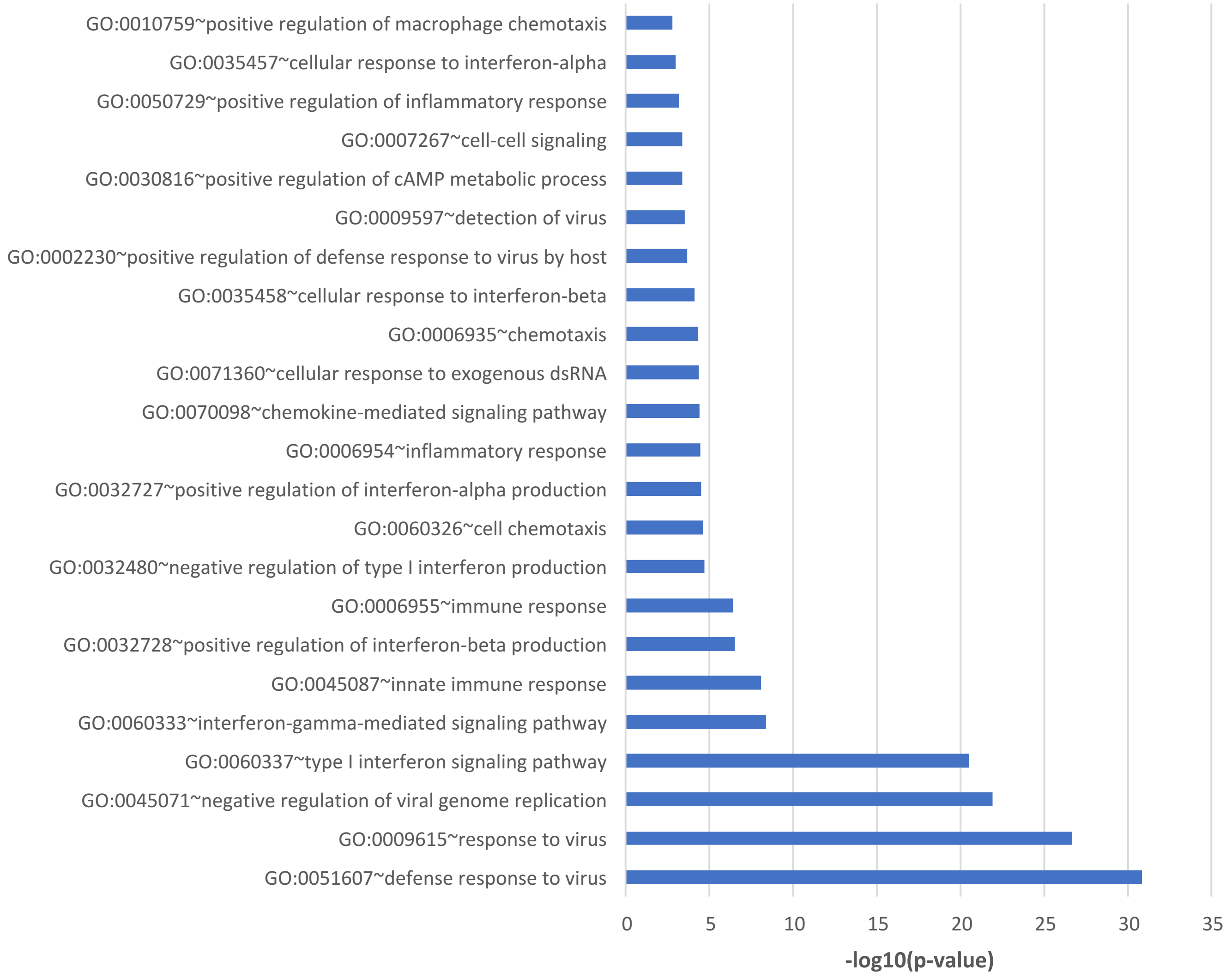


**Figure S4. Genome-Wide Responses to dsDNA** Gene ontology analysis (GOTERM_BP_DIRECT) of genes upregulated more than log(2) in ns sIRNA treated dsVacV-70 transfected cells compared to ns siRNA treated cells transfected with no DNA was performed using DAVID Functional Annotation Clustering Tool.

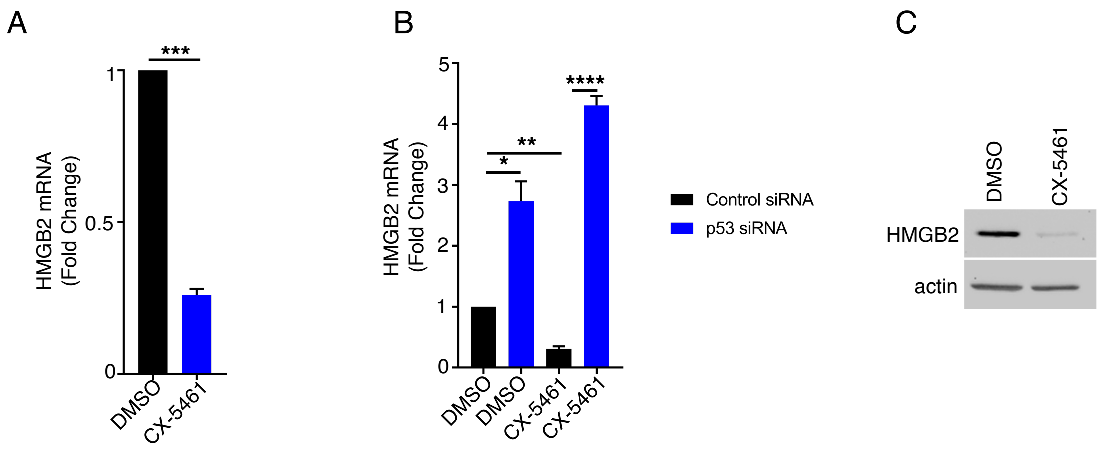


**<u>Figure S5.</u> Inhibiting RNA polymerase I reduces HMGB2 abundance in a p53-dependent manner.** (a) NHDFs were treated with DMSO or 1 mM CX-5461. After 3d, total RNA was isolated and RT-qPCR performed using primers specific for HMGB2 mRNA. Error bars indicate SEM. ***, P≤0.001; Students t-test.(b) NHDFs transfected with the indicated siRNA were treated as in (a).*, P<0.05; **, P<0.01; ****, P<0.0001. (c) As in A except total protein was collected and analyzed by immunoblotting using the indicated antibodies.

| Primer | Sequence (5' - 3') |
|---|---|
| 45S Forward | CGGGTTATTGCTGACACGC |
| 45S Reverse | CAACCTCTCCAGCGACAGG |
| CDCA8 Forward | AGTGGAAATACGAATCAAGC |
| CDCA8 Reverse | TCTCGATGTTGTAGAGGTTATC |
| CDK1 Forward | ACCTATGGAGTTGTGTATAAGG |
| CDK1 Reverse | GACTGACTATATTTGGATGACG |
| CENPF Forward | ACTCACATCAGTAAAGCAAC |
| CENPF Reverse | TCATTCTCCTTGATCTGACTC |
| DDX58 Forward | GGTATAGAGTTACAGGCATTTC |
| DDX58 Reverse | TTGTTTACTAGTGTTGTGGC |
| HMGB2 Forward | GAAAGCAGCCTAAGCTAAAGG |
| HMGB2 Reverse | TTCATCTTCATCCTCTTCCTC |
| IFIH1 Forward | GATTAAGTGGTGATACCCAAC |
| IFIH1 Reverse | GTCTGACAATTGAACACCAG |
| IFIT2 Forward | ACCATGAGTGAGAACAATAAG |
| IFIT2 Reverse | TTAGATAGGCCAGTAGGTTG |
| IFIT3 Forward | ATGAGTGAGGTCACCAAG |
| IFIT3 Reverse | CCTTGAATAAGTTCCAGGTG |
| IFNB1 Forward | GAAAGAAGATTTCACCAGGG |
| IFNB1 Reverse | CCTTCAGGTAATGCAGAATC |
| ISG15 Forward | AGATCACCCAGAAGATCG |
| ISG15 Reverse | TGTTATTCCTCACCAGGATG |
| OAS3 Forward | AGTGTACCAAGATCTCCAAG |
| OAS3 Reverse | ATGGTCCAGTAGATACAGAG |
| pHrP2-BH Forward | ACAAGCTTATTCGCCATTCAGGCTGCGC |
| pHrP2-BH Reverse | TATCTAGACTGGCCGTCGTTTTACAAG |
| RSAD2 Forward | GCTCTAAGAGAAGCAGAAAG |
| RSAD2 Reverse | CATCTTCTGGTTAGATTCAGG |
| TK1 Forward | AAAAGCACAGAGTTGATGAG |
| TK1 Reverse | GAGTGTCTTTGGCATACTTG |
| UBA7 Forward | GGAGACACAACAACTTTCTC |
| UBA7 Reverse | CTTATGTCTCACAGTCTTGG |

**Table S1.** List of primers sequences used for RT-qPCR in this study.

| siRNA | Sequence (5' - 3') |
|---|---|
| TIF-IA #1 | GCCUCAUUGCCUACCAGGA[dT][dT] |
| TIF-IA #2 | CUAGAAUUCCGUUCUUCUA[dT][dT] |
| HMGB2 #1 | CAUCUGCCUUCUUCCUGUU[dT][dT] |
| HMGB2 #2 | CCGUCAAUUUCGCGGAAUU[dT][dT] |
| UBF #1 | GAAGUUCCGUACAUUGACA[dT][dT] |
| UBF #2 | CCAUGUUCAUCUUCUCGGA[dT][dT] |
| TFIIIA #1 | CACUAGGCAUGCUGUUGUA[dT][dT] |
| TFIIIA #2 | AACAUUUGAUUCCUUAUCA[dT][dT] |
| BOP1 | UCGUGCUGAAGUCAACAGA[dT][dT] |

**Table S2.** List of siRNA sequences used in this study.

**<u>Table S3.</u>** List of genes induced by dsDNA in NHDFs treated with non-silencing siRNA, TIF-IA siRNA, or UBF siRNA *(separate excel file)*

**<u>Table S4</u>**. List of genes regulated by TIF-IA depletion in NHDFs. *(separate excel file)*
